# Molecular profiling of frontal and occipital subcortical white matter hyperintensities in Alzheimer’s disease

**DOI:** 10.1101/2024.06.13.598845

**Authors:** Sulochan Malla, Annie G. Bryant, Rojashree Jayakumar, Benjamin Woost, Nina Wolf, Andrew Li, Sudeshna Das, Susanne J. van Veluw, Rachel E. Bennett

## Abstract

White matter hyperintensities (WMHs) are commonly detected on T2-weighted magnetic resonance imaging (MRI) scans, occurring in both typical aging and Alzheimer’s disease. Despite their frequent appearance and their association with cognitive decline, the molecular factors contributing to WMHs remain unclear. In this study, we investigated the transcriptomic profiles of two commonly affected brain regions with coincident AD pathology—frontal subcortical white matter (frontal-WM) and occipital subcortical white matter (occipital-WM)—and compared with age-matched healthy controls. Through RNA-sequencing in frontal- and occipital-WM bulk tissues, we identified an upregulation of genes associated with brain vasculature function in AD white matter. To further elucidate vasculature-specific transcriptomic features, we performed RNA-seq analysis on blood vessels isolated from these white matter regions, which revealed an upregulation of genes related to protein folding pathways. Finally, comparing gene expression profiles between AD individuals with high-versus low-WMH burden showed an increased expression of pathways associated with immune function. Taken together, our study characterizes the diverse molecular profiles of white matter changes in AD compared to normal aging and provides new mechanistic insights processes underlying AD-related WMHs.

## Introduction

White matter hyperintensities (WMHs) are imaging abnormalities observed on T2-weighted MRI scans ^1^. These abnormalities increase with age, with an estimated 90% of individuals over 65 years of age exhibiting such lesions ^2,3^. WMHs may manifest across various regions of the brain, particularly in periventricular, deep frontal subcortical, and parieto-occipital subcortical white matter (WM) areas ^4^. Increased WMH volume is associated with cognitive deterioration and the onset of neurological conditions like Alzheimer’s disease (AD) ^5^. Moreover, WMH volume is predictive of both the development of mild cognitive impairment and the rate of cognitive decline ^4,6-9^.

Despite these clear associations with cognitive decline, the underlying cause(s) of these WM abnormalities remain somewhat unclear ^10^. Factors contributing to WMH formation may include chronic cerebral hypoperfusion ^11,12^, inflammation ^13^, microglial and endothelial cell activation ^14^, blood brain barrier dysfunction ^15^, myelin degeneration, and axonal loss ^16^. Genome-wide association studies indicate that genes regulating blood pressure ^17^ along with other risk factors such as diabetes, hypercholesterolemia, smoking, carotid artery disease, atrial fibrillation, and heart failure ^18^ are associated with WMHs. Considering this large vascular component, WMHs are generally considered as indicators of small vessel disease ^4,17,19-21^. On the other hand, neuropathological examinations comparing frontal and parietal WM lesions in aging and AD indicate that they may be caused by different AD-related mechanisms, such as the deposition of hyperphosphorylated tau and amyloid beta (Aβ) ^3,22-26^.

Altogether, questions remain regarding whether WMHs in AD exhibit unique features compared to normal aging, and whether similar processes underlie WMHs in different brain areas. To address these key questions, we performed RNA-sequencing (RNA-seq) analysis of bulk tissues and blood vessels isolated from the WM of AD cases and age-matched normal controls. Through detailed transcriptomic characterization of WMH-susceptible regions, this study provides insights into the biological underpinnings of these lesions. It provides new evidence that, despite appearing similar by MRI, AD-related WM changes are distinct from normal aging and the development of WMHs reflect a wide array of biological processes including vasculature and protein folding functions.

## Methods

### Human donor tissue collection

All brain tissues included in this study were obtained from the Massachusetts Alzheimer’s Disease Research Center (MADRC) with the consent of the patients or their families and approval from the Mass General Brigham Institutional Review Board (IRB; 1999P009556). Donor tissues were assessed by a neuropathologist and classified as Alzheimer’s disease (AD) or normal aging controls following the NIA-AA guidelines ^27^. Normal aging control cases (n = 5, 3F/2M) were defined as donors with a Braak neurofibrillary tangle (NFT) score of 0/I/II and no history of dementia, whereas AD cases (n = 9, 6F/3M) had extensive tau pathology in the neocortex (Braak V/VI) and were selected based on the availability of *in-vivo* MRI. Some participants in both the control and AD groups exhibited cerebral amyloid angiopathy (CAA) and cerebrovascular disease (CVD) upon neuropathological evaluation (MADRC). For instance, among the 5 control participants, 4 had CVD and 1 had CAA whereas among AD participants, there were 7 with CVD and 6 with CAA. A summary of donor tissues used in this study is presented in **Table 1**.

**Table 1.** Participants demographic information.

| Neuropath group | Sex | Age | Braak Stage | Thal Stage | *In-vivo* WMH volume (mm3) | WMH burden | Death-scan interval (years) | CAA | CVD |
| --- | --- | --- | --- | --- | --- | --- | --- | --- | --- |
| Control-1 | F | 81 | I |  | 0 | NA | NA | Yes | Yes |
| Control-2 | F | 79 | II |  | 0 | NA | NA | No | No |
| Control-3 | M | 89 | II |  | 0 | NA | NA | No | Yes |
| Control-4 | M | >=90 | I |  | 0 | NA | NA | No | Yes |
| Control-5 | F | 89 | I |  | 0 | NA | NA | No | Yes |
| AD-1 | F | 84 | V | 5 | 104 | Low |  | Yes | No |
| AD-2 | M | 84 | V | 5 | 2258 | Low | 2.28 | No | Yes |
| AD-3 | F | 64 | VI | 4 | 2754 | Low | 4.87 | Yes | Yes |
| AD-4 | F | 72 | VI | 5 | 6469 | Low | 1.19 | No | Yes |
| AD-5 | M | 81 | VI | 5 | 11013 | Low | 2.39 | Yes | Yes |
| AD-6 | F | 78 | V | 5 | 23654 | High | 0.69 | Yes | Yes |
| AD-7 | M | >=90 | V | 5 | 37387 | High | 4.87 | Yes | Yes |
| AD-8 | F | 85 | V | 4 | 40529 | High | 3.31 | No | No |
| AD-9 | F | 86 | VI | 4 | 62279 | High | 2.37 | Yes | Yes |

### *In Vivo* MRI image acquisition and processing

Fluid-attenuated inversion recovery (FLAIR) and magnetization-prepared rapid acquisition with gradient echo (MPRAGE) images were collected using a 3T scanner (MAGNETOM Trio, Siemens Healthineers) under existing clinical research protocols approved by the Mass General Brigham IRB and shared via the Neuroimaging and Biomarker Core of the MADRC. FLAIR Images were collected using the following parameters: TE = 303 ms, TI = 2200 ms, TR = 6000 ms with a resolution of 1.0 x 1.0 x 1.0 mm^3^ . MPRAGE used the following parameters: TE = 1.61 ms, TI = 1100, TR = 2530 ms with a resolution of 1.0 x 1.0 x 1.0 mm^3^. FLAIR and MPRAGE DICOM series were unpacked and converted to NIFTI volumes with dcmunpack from FreeSurfer (v7.1.1; https://surfer.nmr.mgh.harvard.edu/). We utilized the sequence adaptive multimodal sequencing (samseg) module from FreeSurfer ^28,29^ with the lesion flag for automatic and unsupervised segmentation of WMH from the native-space FLAIR and MPRAGE volumes. Volumetric segmentation and cortical reconstruction were performed using the standard recon-all pipeline from FreeSurfer (v7.1.1). The FLAIR and samseg-derived WMH volumes were aligned and registered to the FreeSurfer-processed MPRAGE using bbregister from FreeSurfer. WMH volume statistics were derived using the mri_segstats function from FreeSurfer after binarizing the WMH segmentation and co-registering to the FreeSurfer-processed MPRAGE using mri_vol2vol with the .lta file generated from bbregister.

### *Ex vivo* MRI imaging and quantification

Archived formalin-fixed coronal brain slabs (stored for up to 8 years) were rinsed for 24 hours in PBS and then incubated overnight in fresh neutral buffered 10% formalin (Sigma Aldrich, catalog # HT501128). Two slabs from each donor were selected for imaging corresponding to slices nearest the frontal and occipital horn of the lateral ventricles. Slices were stacked in neutral buffered 10% formalin, gently weighted and secured with parafilm (to eliminate bubbles and minimize tissue shifting), and imaged using a 7T MRI scanner (MAGNETOM Trio, Siemens Healthineers) equipped with a 32-channel head coil. A turbo-spin echo (TSE) sequence was acquired (TE = 50 ms, TR = 1000 ms) with a resolution of 0.6 x 0.6 x 0.6 mm^3^ (total scan time 0.5 hours). MRI files were imported to Image-J and regions of interest (ROIs) were manually drawn to isolate each slab from the stack. These images were subsequently used to guide dissection of white matter regions based on brightness relative to the surrounding tissue. Dissected tissues were then embedded in paraffin for downstream histology and immunohistochemistry.

### Histology

Paraffin-embedded frontal-WM and occipital-WM tissues from AD and adjacent control tissues were sectioned on a microtome at 7 µm, and sections were mounted on Superfrost slides as described previously ^30^. Slides containing paraffin-embedded tissue sections were deparaffinized and rehydrated in a sequence of solutions, including (5 minutes each) 100% xylene, 100% ethanol (EtOH), 95% EtOH, 70% EtOH, and distilled H_2_O. For Luxol Fast Blue staining, sections were then stained overnight at 60°C using 0.1 % Luxol Fast Blue (Acros Organics, catalog # AC212170250) in 95% EtOH and 0.05% acetic acid. After staining, sections were rinsed in 95% EtOH and distilled water. Differentiation was carried out by immersing the sections in Lithium Carbonate solution (0.05%; Sigma Aldrich, catalog # L-3876) for 1 minute. Subsequently, sections underwent dehydration with an ascending EtOH series followed by xylene and were coverslipped with Eukitt Quick-Hardening Medium (Sigma, catalog # 03989).

For the validation of targets identified by transcriptomic analysis, tissue sections were deparaffinized in 100% xylene (two changes at 5 min each) followed by rehydration in descending concentrations of EtOH (two changes of 100% EtOH for 5 mins followed by 95%, and 80% EtOH for 1 min). Epitope retrieval was conducted in citrate buffer (pH 6.0, containing 0.05% Tween-20) using microwave heating at 95°C for 20 min (20% power). Sections were then blocked with 5% BSA in TBS containing 0.25% Triton-X for 1 h at room temperature and then incubated with primary antibodies overnight at 4°C. Primary antibodies used in this study targeted HSPA6 (Santa Cruz, catalog # sc-374589), HSP90AA1 (Invitrogen, catalog # PA3-013), and GLUT1 (EMD Millipore, catalog # 07-1401). The following day, sections were rinsed 3 times with TBS and then incubated in Alexa dye-conjugated secondary antibodies in TBS containing 0.25% Triton-X for 1 h at room temperature. The sections were rinsed 3 times with TBS and Trueblack Lipofuscin Autofluorescence Quencher (Biotum, catalog # 23014) was applied for 30 s to quench autofluorescence. Sections were rinsed with PBS and then coverslipped with Fluoromount G with DAPI. For 3,3’-diaminobenzidine (DAB) staining, sections were also pre-treated with 3% hydrogen peroxide (10 min). Following primary antibody incubation, biotinylated secondary antibodies were added for 1 h, then incubated with ABC solution (streptavidin-HRP; Vector Laboratories, catalog # PK-6100) for 1 h and finally, incubated with DAB (Sigma Aldrich, catalog # D5905) in TBS containing nickel chloride for 15 min. DAB-labeled sections were subsequently rinsed in TBS, dehydrated and coverslipped with Eukitt. All images were obtained using an Olympus VS120 automated slide scanner and a 20x lens.

### Isolation of blood vessels from freshly frozen brain tissue

Regions corresponding to subcortical white matter near the frontal and occipital horns of the lateral ventricles were dissected from frozen tissue slabs. These regions were selected *a priori* due to their susceptibility to developing WMHs with aging and AD ^10,26,31^. A 25 mg piece of each region was reserved for bulk RNA sequencing. Blood vessels were isolated from 200-300 mg of frozen frontal- and occipital-WM human tissue from both the control and AD donors following the method previously described ^32,33^. Briefly, tissues were minced in 2 mm sections using a razor blade in ice-cold B1 buffer (Hanks Balanced Salt Solution with 10 mM HEPES, pH 7; Thermo Fisher Scientific). Subsequently, samples were manually homogenized using a 2 ml dounce homogenizer (12 strokes). The resulting homogenized mixture was transferred into a conical tube filled with 20 ml of B1 buffer and centrifuged at 2,000 g for 10 min at 4^0^C. The supernatant was discarded to obtain the pellet, which was then suspended in 20 ml of B2 buffer (B1 buffer with 18% dextran, Sigma-Aldrich) by vigorously mixing for 1 min to separate out myelin. Following this, samples were centrifuged at 4,400 g for 15 min at 4^0^C to remove the myelin layer, after which the pellet was resuspended in 1 ml of B3 buffer (B1 buffer with 1% Bovine Serum Albumin, Sigma-Aldrich). Homogenates were then filtered in a 20 mm mesh (Millipore) pre-equilibrated with 5 ml of ice-cold B3 solution. Brain blood vessels were rinsed with 15 ml of ice-cold B3 solution, detached from the filters by immersing them in 20 ml of B3 ice-cold solution, and centrifuged at 2,000 g for 5 min at 4^0^C. Finally, the pellet was resuspended in 1 ml of ice-cold B1 solution and re-centrifuged at 2,000 g for 5 min at 4^0^C. The supernatant was discarded to obtain the blood vessel pellet, each of which was stored at -80^0^C until further use.

### Bulk RNA-sequencing, analysis, and quality control

Frozen isolated blood vessels or 25 mg bulk brain tissue were placed in buffer RLT (Qiagen) and sonicated with 20 pulses at 10% power. The resulting homogenates were centrifuged at 13,000 rotations per minute (rpm) for 3 min and RNA in the supernatant was purified using the RNase Mini Kit (Qiagen). A NanoDrop spectrophotometer was used to measure the concentration of RNA in each sample. All samples were diluted to 20 ng/µl in RNase free water and sent to Discovery Life Sciences Genomics (Huntsville, AL) for RNA sequencing (RNA-seq). We performed transcript-level abundance quantification from the FASTQ sequencing files using Salmon (v1.5.2) in mapping-based mode ^34^. Briefly, the primary assembly human genome (GRCh38) and full human transcriptome were acquired from GENCODE release 36. The genome was appended to the transcriptome to create a combined “gentrome” file for indexing using salmon’s index command with the gencode flag. Each sample’s paired-end FASTQ reads were then quantified against the full decoy-aware transcriptome using salmon’s *quant* function with –*validate Mappings* and –*gc Bias* flags. We set -l A so Salmon would automatically determine the library type. The Salmon transcript quantification files (quant.sf.gz) were imported into R using the *tximport* package ^34,35^ with a custom constructed TxDb object created from the GENCODE GRCh38 annotation .gtf file using the *makeTxDbFromGFF* function.

### Analysis of differentially expressed genes (DEGs)

Differential gene expression analysis was performed using *DESeq2*, which estimates the mean-variance relationship and log-based fold changes in the count data using the negative binomial distribution ^36^. We constructed the *DESeqDataSet* object using the transcript abundance files and the *tximport* function. For both the AD with WMH (AD) versus control and the high-versus low-WMH AD analyses, we partitioned the data into four groups before running DESeq: (1) frontal-WM bulk tissue; (2) frontal-WM blood vessels; (3) occipital-WM bulk tissue; and (4) occipital-WM blood vessels (BV). We extracted the results from each DESeq analysis using the *results* function, with independent filtering set to FALSE. We considered a gene to be differentially expressed in either AD versus control or high-versus low-WMH comparisons if the Benjamini Hochberg (BH)-adjusted p < 0.05, and magnitude of the log2 fold change (log2FC) > 0.5. The top 30 DEGs per analysis were visualized in a volcano plot by first performing a variance stabilizing transformation (*vst* function) in *DESeq2* and plotting the transformed values.

### Metascape and synapse functional enrichment analysis

Gene ontology (GO) annotation and functional enrichment analysis of DEGs was carried out using Metascape (https://metascape.org) with default settings as previously described ^37^. The Metascape multi-gene-list meta-analysis tool was used to compare common functional pathways among DEGs in AD versus control and high-versus low-WMH burden for both brain regions. For comparative functional gene enrichment annotation obtained from Metascape, DEGs with specified cut off value were submitted to ShinyGO (http://bioinformatics.sdstate.edu/) ^38^. To investigate dysregulated synaptic organization, we submitted DEGs to synaptic ontology database SynGO (https://www.syngoportal.org/) ^39^. The brain expressed background gene set was used to identify the enriched synaptic components.

### Statistical analysis

Individual data points are shown in bar plots, where bar height indicates mean and error bars show ± standard deviations. The volume of *in-vivo* WMH in low-versus high-WMH samples was compared using Student’s t-test in GraphPad Prism (*p < 0.05, **p < 0.01, ***p < 0.001). All other analysis was performed in R as indicated above.

## Results

### Characterization of WMH in control and AD donor tissues

We first selected n=9 AD donors who had undergone T2-weighted MRI scans during life within 5 years prior to death and matched these to n=5 normal aging controls. After review of the *in-vivo* MR images (**Sup. Fig. 1A, B**), we dissected frozen tissues from frontal- and occipital-WM for RNA-seq, as these two areas are commonly affected by WMHs (**Fig. 1A**). In a subset for which formalin-fixed tissues were also available (n = 8), we performed *ex-vivo* MRI of the contralateral hemisphere and observed that WM changes were similar across the normal aging controls and AD donors (**Fig. 1B**). However, we note that the greatest burden of WMH was observed in 2 out of 5 AD cases in these *ex-vivo* images, corresponding to what was observed *in-vivo*. From these formalin-fixed tissues, we dissected additional WM regions and assessed WM integrity by Luxol Fast Blue (LFB) (**Fig. 1C, D**). No significant difference was observed in the percent area of LFB staining within WM, indicating similar myelin status in AD and control samples (**Fig. 1E**). In sum, the integrity of WM was overall similar between AD and normal aging control tissues.

**Figure 1.**
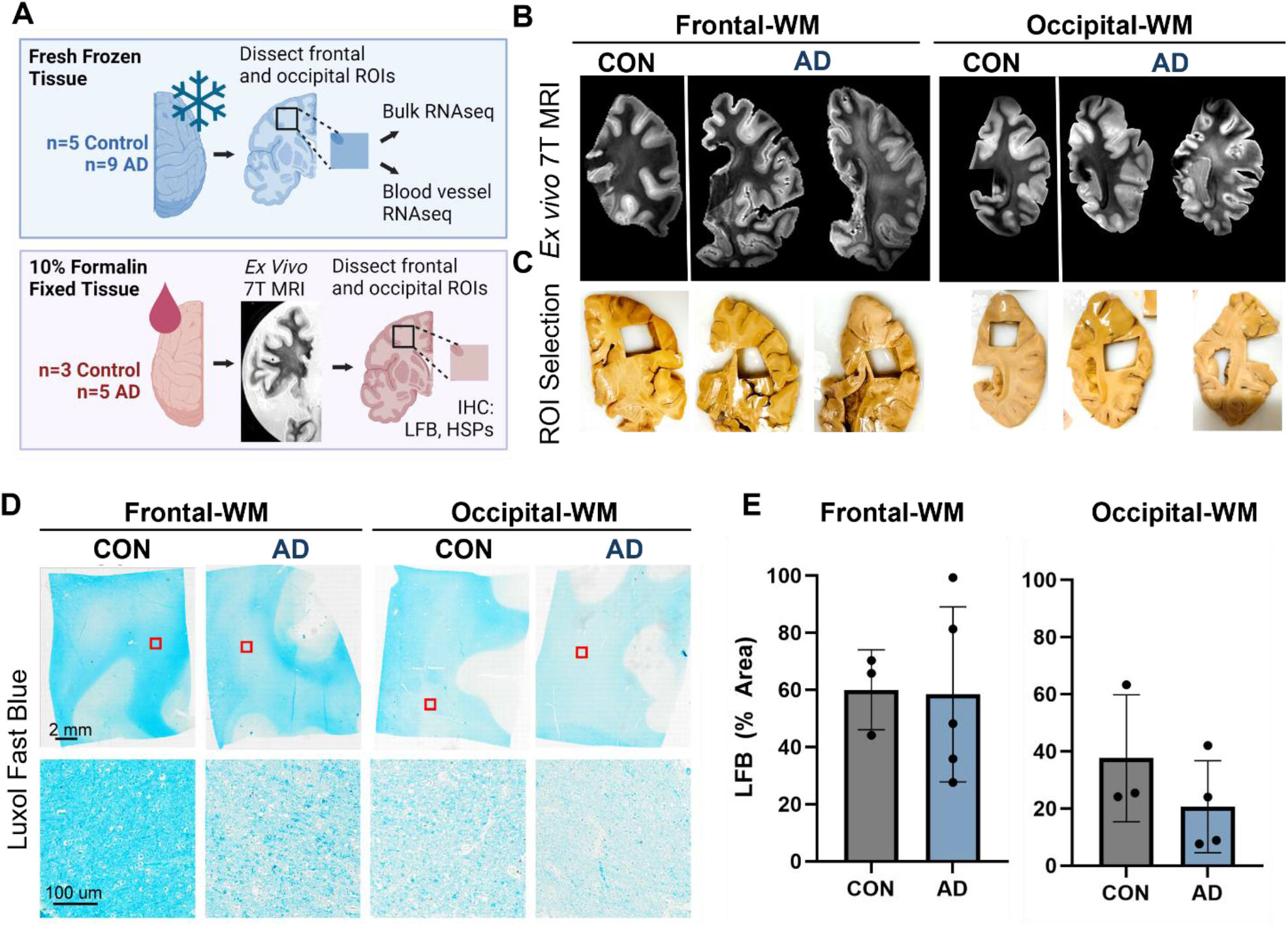
Experimental outline and characterization of WMH in control and AD cases. **A)** Schematic representation of experimental outline and conditions. Tissues were obtained from the Massachusetts ADRC. One hemisphere of the brain was fixed in 10% formalin and was used for *ex vivo* 7T MRI and histopathology. The other hemisphere was freshly frozen, and tissues were used for RNA-seq from the bulk tissues and isolated blood vessels. Not all donors used for RNA-seq had the alternate hemisphere and corresponding brain regions available for histology. **B)** Examples of *ex vivo* T2-weighted TSE MRI scan showing the WMH in AD in frontal- and occipital-WM. **C)** Dissection of region of interest (ROI) showing WMH in the frontal- and occipital-WM as indicated by the *ex vivo* MRI images in (B). **D)** Luxol Fast Blue (LFB) staining of the control and AD samples in frontal- and occipital-WM. **E)** The percentage area of LFB staining is shown in control and AD for frontal- and occipital-WM.

### Alterations in brain vasculature pathways are underlying features of AD WM

We next sought to examine transcriptomic alterations unique to AD WM compared to controls. In frontal-WM bulk tissues, RNA-seq identified 2198 DEGs, of which 1033 were significantly upregulated (log2FC > 0.5, padj. < 0.05) in AD versus controls. Notable among upregulated genes are *RNA5S1, LRRC71, CCDC194, IL5RA, DUX4, CTAGE4, TTR* and *MLKL* (**Fig. 2A, Sup. Table 1)** and increased transcription of pseudogenes such as *NSFP1, MT1JP*, and *DNAJB1P1*. To examine the biological roles of the significantly upregulated genes, we performed functional gene enrichment analysis, which highlighted biological processes including DNA damage response, tube morphogenesis and regulation of Hippo-signaling (**Fig. 2B, Sup. Table 2**). Specifically, DNA damage included subclasses such as DNA metabolic process and DNA double strand break repair while tube morphogenesis encompasses blood vessel development, blood vessel morphogenesis, and angiogenesis (**Sup. Table 2)**.

**Figure 2.**
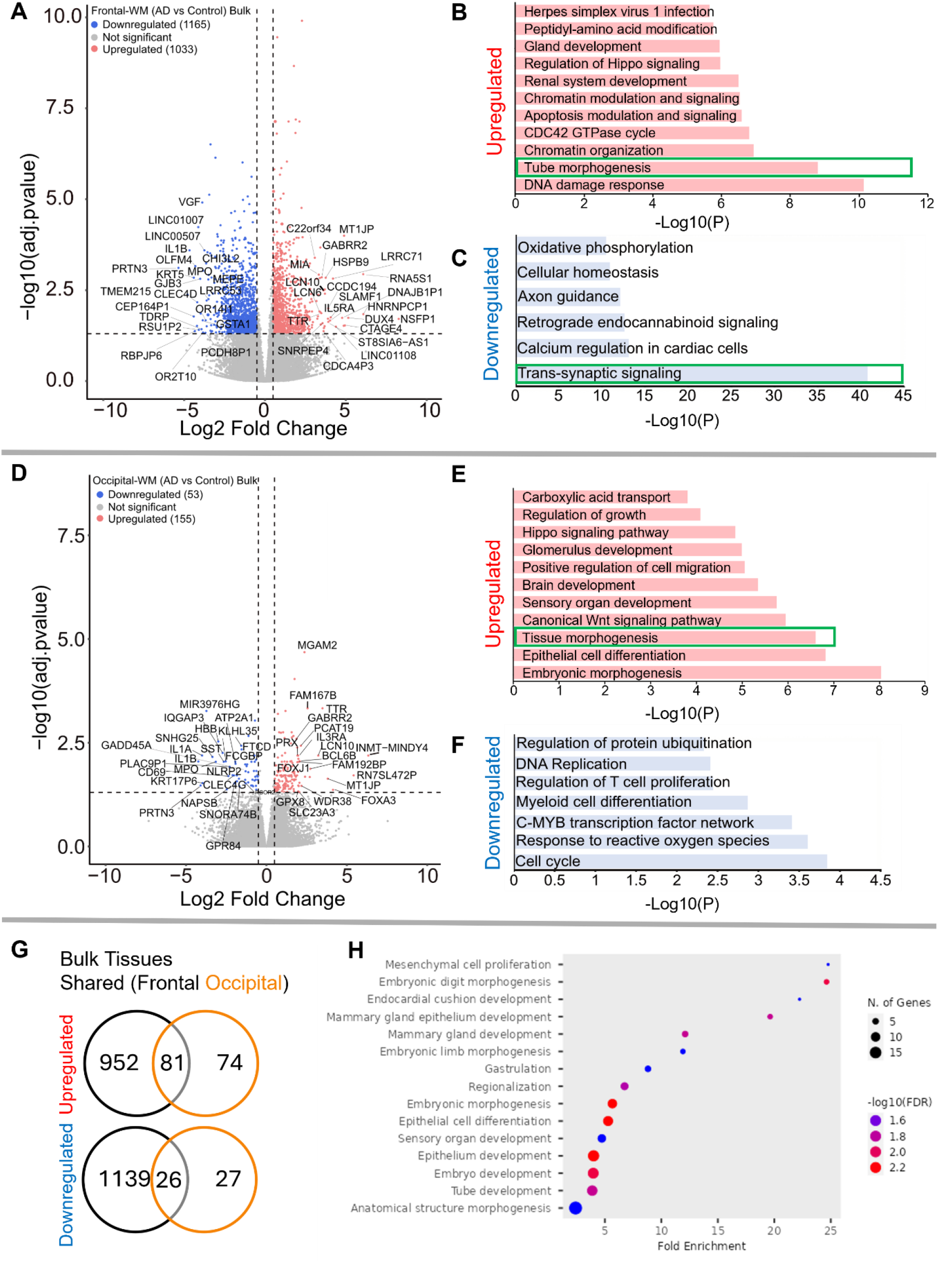
Gene expression changes in frontal-and occipital-WM bulk tissues. **A** and **D**) Volcano plot showing differential gene expression in **(A**) Frontal-WM bulk tissues and **(D)** Occipital-WM bulk tissues. The color of dots indicates the direction of significantly altered genes, with red indicating upregulation and blue indicating downregulation. **B-C & E-F)** Bar plot showing the enriched biological processes (GO/KEGG terms, canonical pathways) generated by Metascape for (**B & E**) upregulated genes, and (**C & F**) downregulated genes. The enriched pathways associated with brain vasculature in figure E and F and synaptic functions in figure C are marked by green boxes. **G)** Venn diagram showing unique and shared DEGs in the frontal- and occipital-WM bulk tissues for upregulated genes (upper panel), and downregulated genes (lower panel). **H)** Biological processes associated with upregulated genes common to frontal- and occipital-WM bulk tissues, generated by ShinyGO. The enriched pathways associated with brain vasculature functions are marked by yellow boxes.

Conversely, the 1165 downregulated transcripts (log2FC < -0.5, padj. < 0.05) in frontal-WM included *VGF, OLFM4, PRTN3, TMEM215, IL1B, KRT5, CHI3L2, GJB3, MPO*, and *GSTA1* **(Fig. 2A, Sup. Table 1)**. Functional enrichment analysis of these significantly downregulated genes revealed pathways linked to trans-synaptic signaling, axon guidance, cellular homeostasis, and oxidative phosphorylation (**Fig. 2C, Sup. Table 2**). Representative trans-synaptic signaling genes include *SYT1, SYT2, SYT4, SYT5, SYT16, SYTL2, SYTL5, SNAP25, VAMP1, OLFM4*. The enrichment of synaptic functions prompted us to further evaluate the synaptic location and functional GO of downregulated genes using SynGO, an experimentally annotated database for synaptic location and functional GO ^39^. None of the upregulated genes were enriched for synaptic components (**Sup. Fig. 2A-2B**). By comparison, downregulated genes were highly enriched for transcripts known to be related to synaptic location (**Sup. Fig. 2C**) and function (**Sup. Fig. 2D**). Collectively, the differential gene expression observed in frontal-WM bulk tissue primarily indicates alterations in biological processes related to DNA damage, brain vasculature and synaptic function.

Examination of occipital-WM bulk tissues revealed only 208 DEGs between AD and control **(Sup. Table 3**), with 155 upregulated transcripts including the genes *TTR, LCN10, MGAM2, FOXA3, FAM167B* and *WDR38* **(Fig. 2D)**. Biological processes associated with these upregulated genes included epithelial cell differentiation and tissue morphogenesis **(Fig. 2E)**. Interestingly, subclasses within tissue morphogenesis showed many vasculature processes, including tube morphogenesis and regulation of endothelial cell migration (**Sup. Table 4**). Moreover, we also observed enrichment of canonical Wnt-signaling and Hippo signaling pathways, which have been shown to regulate vascular networks and angiogenesis through endothelial cell proliferation and migration ^40^. This observation suggests that, as with frontal WM, vasculature processes are one among many notable altered biological processes in the occipital-WM.

Significantly downregulated genes in occipital-WM included *IL1A, IL1B, SNHG25, PRTN3, MPO* and *SST* **(Fig. 2D)**. Functional enrichment annotation revealed that the downregulated genes in occipital-WM bulk tissues were linked to cell-cycle regulation and response to reactive oxygen species (ROS) (**Fig. 2F**). Of note, none of the occipital-WM DEGs (either up- or down-regulated) were associated with synaptic processes (**Sup. Fig. 2E, F**).

To further compare WM transcriptome in these brain regions, we next examined unique and shared DEGs between frontal- and occipital-WM AD versus control bulk tissues. Although most altered genes in frontal-WM were unique, 52.3% of upregulated (81/155) and 49.0% of downregulated genes (26/53) in occipital-WM overlapped with frontal-WM bulk tissues (**Fig. 2G**). Analysis of these shared upregulated genes using ShinyGO (v. 0.80) confirmed developmental pathways are universally upregulated in AD WM tissue (**Fig. 2H)**. Collectively, our findings from both frontal- and occipital-WM confirm that vasculature alterations are a notable shared feature of AD WM pathology while changes affecting WM synaptic functions are more regionally specific.

### Blood vessels from frontal- and occipital-WM upregulate protein folding genes

Since both frontal- and occipital-WM showed shared gene expression changes related to brain vasculature function, we next investigated gene expression changes within isolated blood vessels. Frontal-WM blood vessels exhibited 690 DEGs. Of these, 568 were upregulated, including *LCN10, KLF15, FOXJ1, CRLF1, CFAP126, CFAP52, MAPK15, CFAP65, FNDC1, GMNC, SPAG17, FAM81B, FAM166C, LHX9, TRDN*, and *OTOS* (**Fig. 3A, Sup. Table 5)**. Notable biological processes associated with these upregulated genes include axoneme assembly, protein folding, chaperone mediated autophagy, and response to metal ions (**Fig. 3B, Sup Table. 6**). Interestingly, protein folding emerged as one of the top altered biological processes after axoneme assembly. We also identified 122 downregulated transcripts in frontal-WM blood vessels, which were associated with multiple biological pathways including ribosome disassembly, RNA splicing, and regulation of extracellular matrix organization (**Fig. 3C)**.

**Figure 3.**
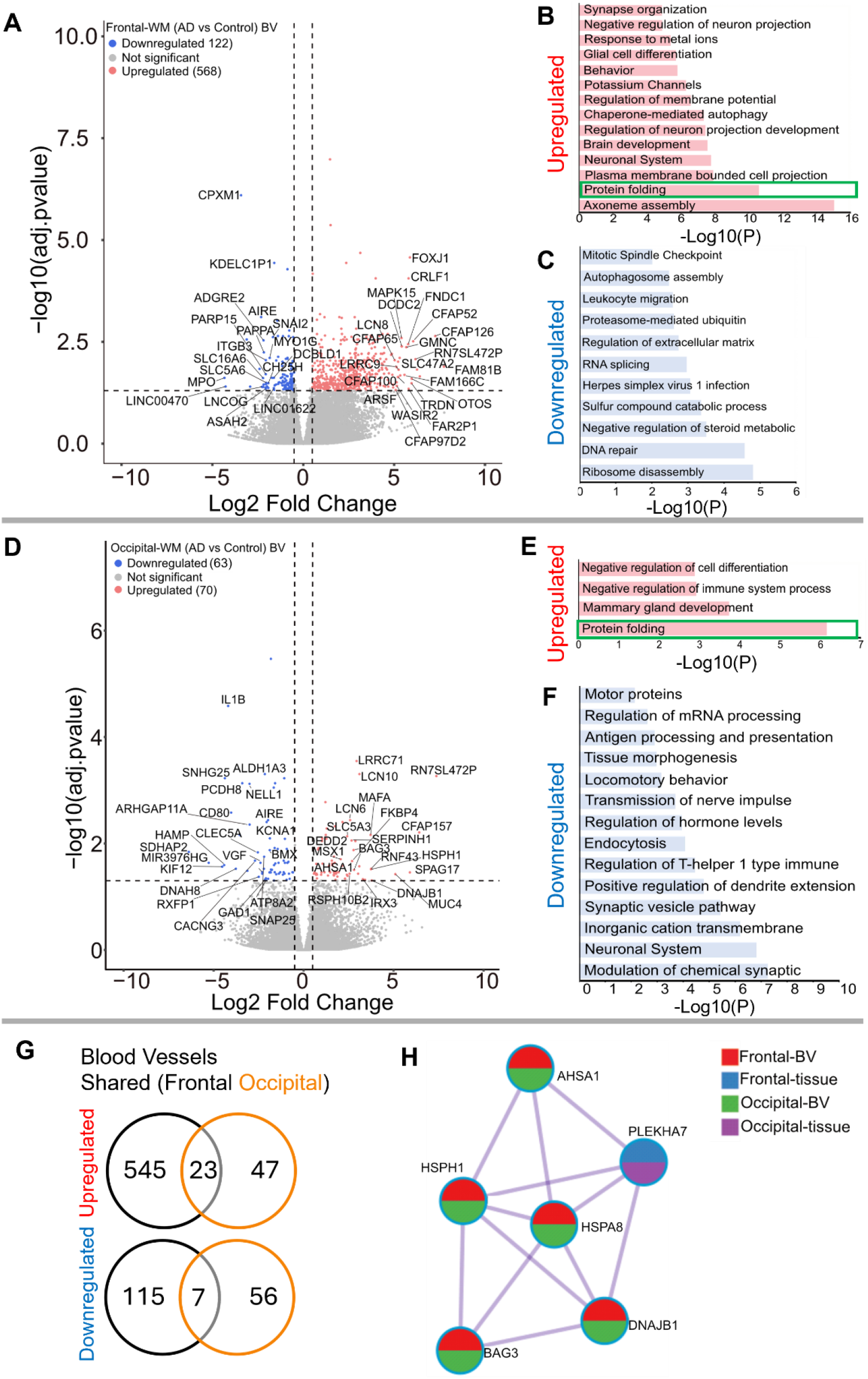
Gene expression changes in isolated blood vessels from frontal-and occipital-WM in AD cases. **A & D**) Volcano plots showing differential gene expression in blood vessels (BV) of (**A**) Frontal-WM and (**D**) Occipital-WM. The color of dots indicates the direction of significantly altered genes, with red indicating upregulation and blue indicating downregulation. **B-C & E-F)** Bar plot showing the enriched biological processes (GO/KEGG terms, canonical pathways) generated by Metascape for (**B & E**) upregulated genes, and (**C & F**) downregulated genes. Enriched pathways associated with protein folding in figure B and E are indicated by green boxes. **G)** Venn diagram showing unique and shared DEGs in the isolated blood vessels from frontal- and occipital-WM for upregulated genes (upper panel) and downregulated genes (lower panel). **H**) Enrichment network visualization for results from the significantly upregulated gene lists in AD versus control, color code represents that HSPs are generally shared among all RNA-seq samples.

Next, we examined if the protein folding pathway observed in frontal-WM blood vessels was also upregulated to occipital-WM blood vessels. Towards this goal, we first explored DEGs in occipital-WM blood vessels which yielded 133 total transcripts. Of these, 70 were upregulated including *LCN10, KLF15, CFAP157, SPAG17, MUC4, HSPH1, FKBP4*, and *HSPA8* (**Fig. 3D, Sup. Table 7**). Functional gene enrichment analysis showed that upregulated genes in occipital-WM blood vessels were most strongly associated with the protein folding pathway (**Fig. 3E, Sup. Table 8**). Last, in occipital-WM blood vessels, we observed 63 downregulated transcripts that were associated with the modulation of inorganic cation transport, endocytosis, and antigen processing (**Fig. 3F**).

While AD blood vessels isolated from frontal- and occipital-WM brain regions shared few up- or down-regulated transcripts (**Fig. 3G**), we noted that at the pathway level, both brain regions were enriched for genes related to protein folding (**Fig. 3B, E)**. This enrichment included chaperone-mediated and de novo-protein folding, heat shock factor (HSF)-dependent transactivation, and protein maturation (**Sup. Tables 6 and 8)**. We also observed more pronounced upregulation of heat shock proteins (HSPs) in blood vessels isolated from both frontal- and occipital-WM blood vessels than in bulk tissues (**Fig. 3H**), which are one of the major components of the protein folding pathway ^41,42^. To confirm the observed changes in the protein folding response of vascular cells, we performed histopathological validation of representative HSPs in sections from frontal-WM of donors used in this study. Consistent with the significant expression of *HSP90AA1* transcripts in blood vessels from AD frontal-WM (log2FC 2.98, padj. = 0.0091, **Sup. Table 5**), HSP90AA1 protein was also expressed in AD frontal-WM blood vessels (**Fig.4A**). We also observed elevated gene expression of *HSPA6* (log2FC 4.83, padj. 0.0066, **Sup. Table 5**) and confirmed protein expression by histology (**Fig. 4B)**. In sum, these observations indicate widespread proteostasis dysfunction in WM vasculature with AD pathology.

**Figure 4.**
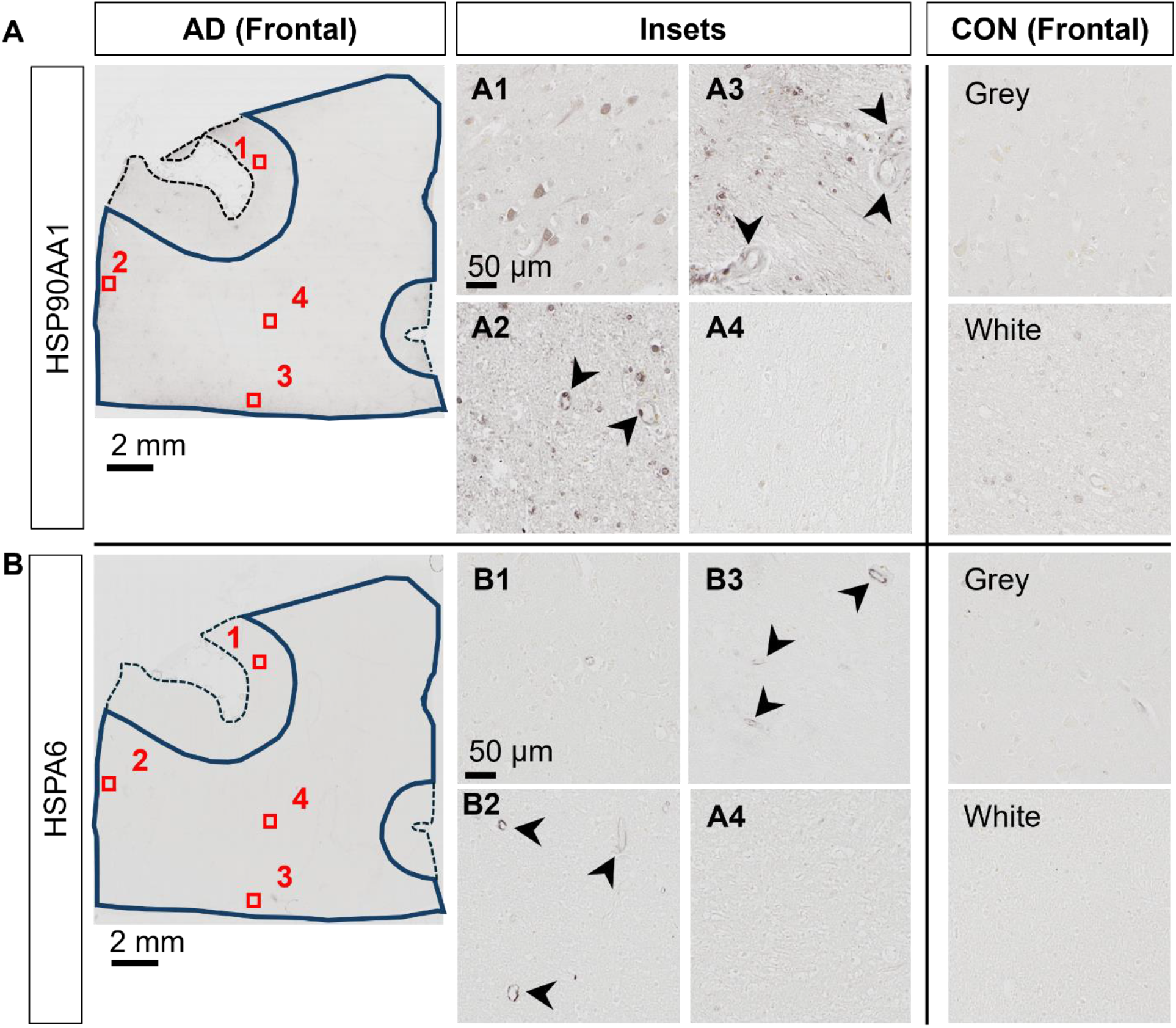
Histopathological validation of heat shock proteins. **A-B**) Immunohistochemistry of frontal-WM tissue section for **A**) HSP90AA1, and **B**) HSPA6 in adjacent tissue sections. Left panels show an AD donor, with closer examination of numbered regions of interest show in insets. Arrowheads highlight vascular cells with HSP-positivity. Right panels show representative control brain white and grey matter areas.

### Analysis of AD donors with high-WMH burden reveals distinct, regional upregulation of genes associated with immune function

Having characterized the transcriptomic differences in the tissues and blood vessels in AD relative to controls, we next explored gene expression differences in AD donors with low- or high-WMH burden. *A priori*, donors were classified into these groups based on *in-vivo* MRI Fazekas scores ^43^ (**Fig. 5 A-D**; low = periventricular WMH ≤2, deep subcortical WMH ≤1; high = periventricular WMH >2, deep subcortical WMH >1). Comparison of total brain WMH volume measured from *in-vivo* MRI revealed a significant difference between groups (p-value = 0.0019; **Fig. 5E**): AD donors in the low-WMH group had WMH volumes of 100 - 11,000 mm^3^ while AD high-WMH donors ranged from 23,654 - 62,279 mm^3^. The average ages for low- and high-WMH AD patients were 77 ± 8.8 (n = 5) and 85 ± 5.4 (n = 4; **Sup. Fig. 1B**), respectively, and both groups had similar AD neuropathology (Braak V-VI and Thal 4-5 stages: **Sup. Fig. 1B**). Although WMH is widely recognized as a feature of aging that increases with age ^2,3,44^, we did not observe a statistically significant correlation between age and WMH volume in our sample (p-value 0.14) despite a positive correlation (Pearson r = 0.54) (**Fig. 5F**). Given their similarities in age and AD pathology status, we next used samples from these donors to probe whether gene expression changes were further enhanced by WMH burden.

**Figure 5.**
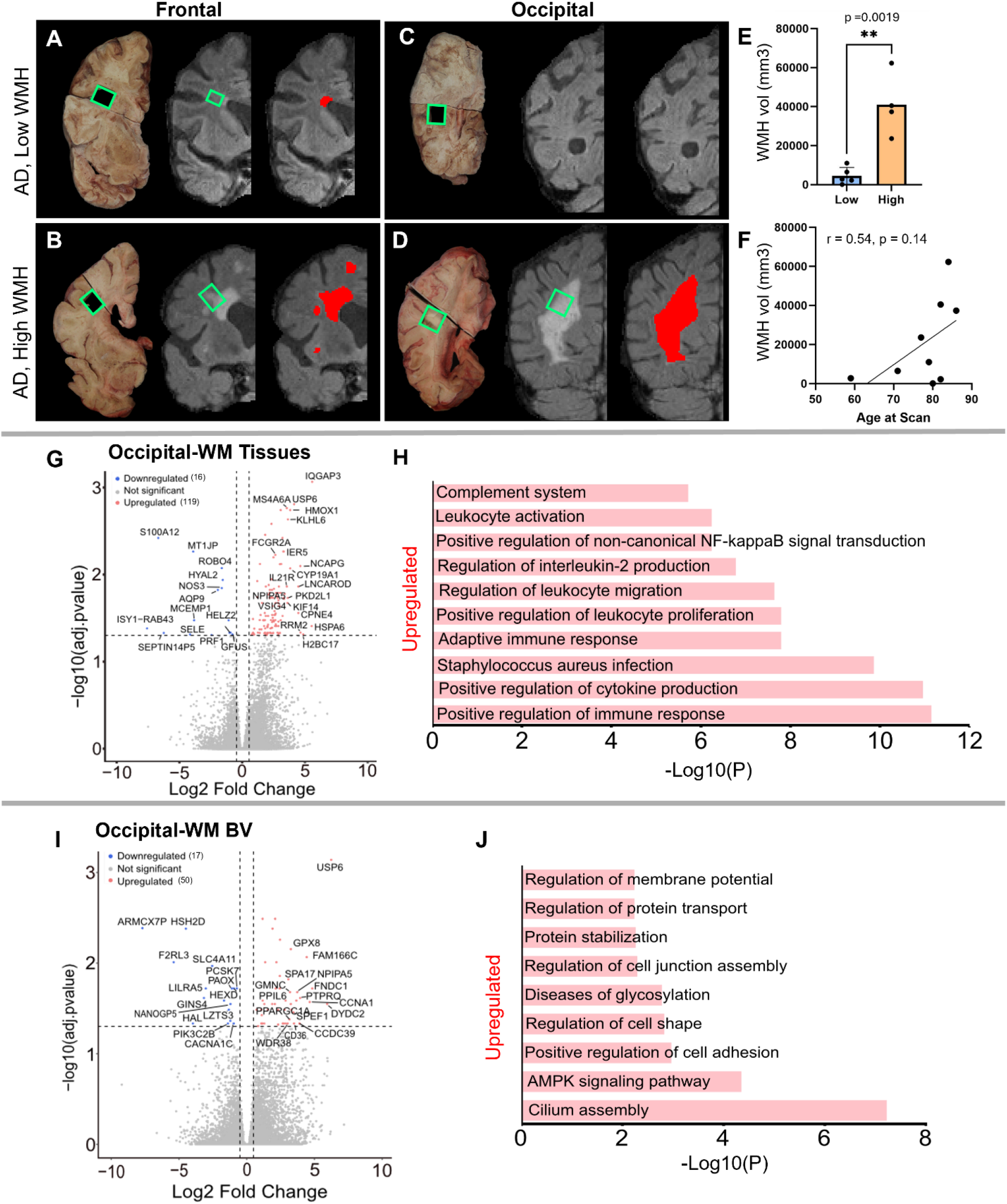
Characterization of low- and high-WMH in frontal- and occipital-WM in AD cases. **A)** Examples of *in vivo* T2-weighted FLAIR MRI scans showing segmented WMH volumes (red pixels) in AD in frontal-WM (**A & B**) and occipital-WM (**C & D**). Tissue sections indicated by green boxes corresponding to the *in vivo* WMH (red and grey pixels) from AD presenting low-WMH (**A & C**) and high-WMH (**B & D**) were dissected. **E)** Comparison of high- and low-WMH samples based on volume of WMH region. **F)** Pearson correlation between the *in vivo* MRI (volume) and age at the time of the scan. **G-I)** Volcano plot showing differential gene expression in low- versus high-WMH in occipital-WM from **G)** bulk tissues, and **I)** blood vessels. **H-J**) Bar plot showing the enriched biological processes (GO/KEGG terms, canonical pathways) generated by Metascape for low- versus high-WMH in occipital-WM from **H)** bulk tissues, and **J)** blood vessels.

Differential gene expression analysis in the high-versus low-WMH groups yielded few transcripts in the frontal-WM after adjusting for multiple comparisons. In the frontal-WM bulk tissue, we found only two DEGs: *POLQ* (log2FC 2.8, padj. 0.01) and *BCAT2* (log2FC -0.6, padj. = 0.0005) (**Sup. Table 9**). DNA polymerase theta (*POLQ*) has been shown to repair double-strand DNA breaks ^45^. Mitochondrial branched-chain amino-acid aminotransferase (*BCAT2*) is confined to vascular endothelial cells, where it triggers dysregulation of glutamate metabolism ^46,47^. In blood vessels isolated from the frontal-WM, only 3 genes were upregulated (*NPIPA5, TPM3P9*, and *CWF19L2*) and two were downregulated (*MT-RNR1* and *PLPPR3)* (**Sup. Table 10)**. This suggests that transcriptional changes in frontal WM are not closely related to WMH burden.

By contrast, occipital-WM exhibited more DEGs in tissue (**Fig. 5G, Sup. Table 11**) and isolated blood vessels (**Sup. Table 12**) with increasing WMH burden. We found 119 upregulated transcripts in occipital-WM bulk tissue including the genes *IQGAP3, HSPA6, H2BC17, RRM2, NCAPG, CPNE4, USP6, CYP19A1, HMOX1, KLHL6, PKD2L1, KIF14, IL21R* (**Sup. Table 11**). Functional gene enrichment analysis showed that these upregulated genes in occipital WM bulk were predominantly associated with immune function encompassing cytokine production, adaptive immune response, leukocyte proliferation and migration, interleukin-2 production and complement system (**Fig. 5H, Sup. Table 13**). In blood vessels from occipital-WM, we found 50 upregulated transcripts including *USP6, DYDC2, FNDC1, CCNA1, FAM166C, PTPRQ, GRIN3B, CCDC39, NPIPA5, CLIC6, CD36* (**Fig. 5I, Sup. Table 12**). These upregulated genes were primarily associated with cilium assembly, AMPK signaling pathway, regulation of cell adhesion, and protein stabilization (**Fig. 5J, Sup. Table 14**). We also found 16 and 17 downregulated transcripts in occipital-WM bulk tissues and blood vessels respectively, which includes *NOS3, HYAL2, AQP9, and SELE* (tissue**; Sup. Table 11**) and *CACNA1X, PAOX, and SLC4A11* (blood vessels; **Sup. Table 12**). In sum, these data indicate that the extent of occipital WMH burden in individuals with AD is associated with greater inflammation, immune signaling, and cell adhesion.

## Discussion

White matter hyperintensities (WMH) are common age-associated neuroradiological changes associated with cognitive decline in Alzheimer’s disease (AD) ^4,17,48-52^. While WMHs are hypothesized to arise from vascular origins, the underlying molecular changes taking place in WMH-prone areas, and whether these differ between AD and normal aging, are not well characterized. In this study we explored disease-related molecular features by RNA-seq comparison with age-matched healthy controls and compared two spatially distant WM regions. We found notable overlap in gene expression programs related to blood vessels and protein folding pathways that are upregulated in WM with disease, but also noted key differences between regions. In particular, comparative analysis between AD groups with high- or low-WMH burden identified upregulation of pathways related to immune function and leukocyte activation particularly in the occipital-WM. Altogether, these data indicate that unique molecular pathways may drive WM changes in AD versus normal aging and even between WMH-susceptible brain regions.

Recent studies have shown that both the location ^53^ and severity ^52^ of WMHs across brain regions are associated with cognitive functions ^54^. Moreover, another study found that posterior-WMHs are linked to AD-related neurodegeneration, while anterior-WMHs are associated with both AD-related neurodegeneration and vascular mechanisms ^55^. These findings underscore the need to understand the biological mechanisms underlying WMHs throughout the brain. In our comparative analysis of two commonly affected WM brain regions, we found far more DEGs in the AD frontal-WM than occipital-WM control bulk tissues. Moreover, we found that genes associated with synapse function were significantly downregulated in AD frontal-but not occipital-WM bulk tissues. Synaptic structures in WM have previously been observed and may be related to modulation of oligodendrocytes and astrocytes, though the significance of their downregulation in AD remains to be characterized ^56,57^. A similar involvement of synaptic input has also been observed in the cortical WM in human temporal lobe epilepsy ^58^, suggesting the involvement of synaptic dysregulation in WM. In sum, this highlights key differences in the molecular changes taking place between frontal- and occipital-WM in AD.

We also found that alterations in pathways associated with tube/tissue morphogenesis, vasculature development, and sprouting angiogenesis were upregulated in both frontal and occipital bulk WM tissues from AD donors. This is in line with the prevailing understanding that WMH changes are associated with vascular dysfunction ^10^. We also found a few downregulated genes common to both brain regions, including *PRTN3, SST*, and *MPO*. Proteinase 3 (*PRTN3)* has been previously reported to regulate endothelial barrier function ^59^ whereas somatostatin (*SST)* has been shown to modulate cortical circuits and cognitive function ^60^. Myeloperoxidase (*MPO)* regulates oxidative stress, suggesting its downregulation may result in elevated ROS levels thus triggering the activation of stress response pathways ^61^. Closer examination of gene expression changes in blood vessels in frontal- and occipital-WM shows upregulated genes are associated with protein folding. The enrichment network visualization of upregulated genes from AD donors indicates that HSPs are generally shared among bulk tissues and blood vessels in both frontal- and occipital-WM brain regions, albeit with higher frequency in blood vessels (**Fig. 3H**). The increased expression of genes associated with HSPs and protein folding pathway are a crucial part of the translational machinery and proteostasis, suggesting that WMH in AD induces localized and wide-spread proteostatic stress in the brain. This finding is consistent with previous reports of upregulated heat shock-related protein folding pathways in AD based single nucleus RNA-sequencing of endothelial cells from multiple grey matter cortical regions ^30^.

Interestingly, while comparison between control and AD donors yielded many DEGs, only two genes that were significantly upregulated across both regions in bulk tissue and blood vessels: lipocalin 10 (*LCN10*) and Krüppel-like factor 15 (*KLF15*). A recent study in mice highlights that LCN10 protects inflammation induced vascular leakage by regulating endothelial barrier integrity and permeability through an LRP2-Ssh1 mediated signaling pathway ^62^. Thus, the upregulation of *LCN10* suggests potential involvement in maintaining the vascular integrity in AD with WMHs. Likewise, *KLF15*, a zinc finger DNA-binding protein belonging to the Krüppel-like family of transcription factors, has also been shown to protect vascular endothelial dysfunction induced by tumor necrosis factor-alpha (TNF-α) in cultured human endothelial cells ^63^. Given this existing literature supporting the role of these genes in blood brain barrier dysfunction, this further supports that vasculature-related processes are key features of AD-related changes.

The transcriptomic analysis of the bulk tissue and blood vessels isolated from high-WMH in occipital-WM revealed that upregulated genes were related to the positive regulation of immune response, regulation of cell activation, positive regulation of cytokine production, and myeloid leukocyte activation. This is in line with a recent study demonstrating a correlation between leukocyte gene expression and WMH progression ^64^ and a recent case report indicating greater microglial burden in hyperintense WM areas compared to adjacent tissue ^65^. Moreover, another group employed epigenomic and integrative-omics data to identify 19 pivotal regulatory genes that influences WMH burden ^66^. These genes were associated with immune function, blood brain barrier, extracellular matrix organization, lipid, and lipoprotein metabolism ^66^. Another transcriptomic study demonstrated an augmented volume of WMH with the increased expression of 39 genes ^67^. Similarly, single-cell RNA sequencing of monocytes isolated from patients with cerebral small vessel disease showed an increased expression of cytokine production and pro-inflammatory markers ^67^. Consistent with our observations, comprehensive analysis of whole blood gene expression identified upregulation of cytokine-cytokine receptor interaction and B-cell receptor signaling ^68^. Our findings are also in line with a recent epigenetics study that demonstrated DNA methylation is linked to WMH formation, particularly influencing blood-brain barrier function and immune response ^69^.Thus, the data presented here and elsewhere indicate that brain vasculature and immune functions are major altered pathways associated with WMH.

While this comprehensive RNA-seq study highlighted potential drivers of AD WMHs, there are limitations to this work that we hope to address with future studies. For one, given a small sample size and variations in gene expression among samples, a larger population-based cohort including diverse ethnic groups^70^ may further substantiate the current findings. Second, our current samples were chosen based on the availability of *in vivo* MRI data. Consistent intervals between T2-FLAIR scans and death are difficult to obtain, and while we only selected cases with scans <5 years prior to tissue collection, we cannot exclude the possibility that some individuals who were classified as having a “low” WMH burden did not subsequently develop additional lesions ^4,64,71,72^. Third, no *in vivo* MRI data was available for the control tissues, so this study did not directly compare WMH versus normal-appearing white matter within the cases.

While we qualitatively examined WMH *ex vivo*, these images were acquired from the hemisphere contralateral to the ones used for RNA-seq per participant; despite our best attempt to match regions, it may not be possible to perform a one-to-one comparison between *in vivo* and *ex vivo* measures. Finally, variables other than WMH burden, such as genetic risk factors not assayed here, may further contribute to gene expression differences between AD donors and WMH burdens. In the future, additional histological examination of key gene targets described in individuals with WMH and AD would help confirm our findings and compare them to other neurological diseases.

In conclusion, our transcriptomic analysis underscores the role of brain vasculature processes and protein folding pathways as key contributors to global WM changes in the AD brain. While these features were shared between brain regions, additional gene expression changes were observed only in frontal-WM indicating separate processes are at work including downregulation of synaptic pathways in disease. Moreover, enhanced upregulation of cytokine production and immune functions were related to greater occipital WMH burden in individuals with AD providing new insights as to how these lesions may contribute to faster cognitive decline in individuals with AD, though follow-up studies are warranted to confirm this finding.

## Data availability

The raw sequencing files will be deposited in SRA upon publication of this dataset. Additional data, including processed files, that support the findings of this study are available from the authors upon reasonable request.

## Acknowledgements

This work was supported by the National Institute on Aging grant R01AG071567 to REB. We thank the donors and their families who have contributed to the Massachusetts Alzheimer’s Disease Research Center, which is supported by the NIH NIA P30AG062421. We additionally would like to thank Tessa R. Connors, Lexi Melloni, and Angelica Gaona for their technical support and assisting with human postmortem tissues used in this study. Finally, we also thank Manolis Maragakis (NIH/NIA) for helpful discussion on this manuscript.

## References

1 Merino, J. G. White matter hyperintensities on magnetic resonance imaging: What is a clinician to do? Mayo Clin. Proc. 94, 380–382 (2019). 10.1016/j.mayocp.2019.01.016

2 de Leeuw, F. E. et al. Prevalence of cerebral white matter lesions in elderly people: a population based magnetic resonance imaging study. The Rotterdam Scan Study. J. Neurol. Neurosurg. Psychiatry 70, 9–14 (2001). 10.1136/jnnp.70.1.9

3 Launer, L. J. et al. Regional variability in the prevalence of cerebral white matter lesions: an MRI study in 9 European countries (CASCADE). Neuroepidemiology 26, 23–29 (2006). 10.1159/000089233

4 Alber, J. et al. White matter hyperintensities in vascular contributions to cognitive impairment and dementia (VCID): Knowledge gaps and opportunities. Alzheimers Dement. (N. Y.) 5, 107–117 (2019). 10.1016/j.trci.2019.02.001

5 Prins, N. D. & Scheltens, P. White matter hyperintensities, cognitive impairment and dementia: an update. Nat. Rev. Neurol. 11, 157–165 (2015). 10.1038/nrneurol.2015.10

6 Kloppenborg, R. P., Nederkoorn, P. J., Geerlings, M. I. & van den Berg, E. Presence and progression of white matter hyperintensities and cognition: a meta-analysis. Neurology 82, 2127–2138 (2014). 10.1212/WNL.0000000000000505

7 Kynast, J. et al. White matter hyperintensities associated with small vessel disease impair social cognition beside attention and memory. J. Cereb. Blood Flow Metab. 38, 996–1009 (2018). 10.1177/0271678X17719380

8 Vergoossen, L. W. M. et al. Interplay of white matter hyperintensities, cerebral networks, and cognitive function in an adult population: Diffusion-tensor imaging in the Maastricht Study. Radiology 298, 384–392 (2021). 10.1148/radiol.2021202634

9 Puzo, C. et al. Independent effects of white matter hyperintensities on cognitive, neuropsychiatric, and functional decline: a longitudinal investigation using the National Alzheimer’s Coordinating Center Uniform Data Set. Alzheimers Res Ther 11, 64 (2019). 10.1186/s13195-019-0521-0

10 Wardlaw, J. M., Valdés Hernández, M. C. & Muñoz-Maniega, S. What are white matter hyperintensities made of? Relevance to vascular cognitive impairment. J. Am. Heart Assoc. 4, 001140 (2015). 10.1161/JAHA.114.001140

11 Tomimoto, H. et al. Chronic cerebral hypoperfusion induces white matter lesions and loss of oligodendroglia with DNA fragmentation in the rat. Acta Neuropathol. 106, 527–534 (2003). 10.1007/s00401-003-0749-3

12 Fernando, M. S. et al. White matter lesions in an unselected cohort of the elderly: molecular pathology suggests origin from chronic hypoperfusion injury. Stroke 37, 1391–1398 (2006). 10.1161/01.STR.0000221308.94473.14

13 Swardfager, W. et al. Peripheral inflammatory markers indicate microstructural damage within periventricular white matter hyperintensities in Alzheimer’s disease: A preliminary report. Alzheimers Dement. (Amst.) 7, 56–60 (2017). 10.1016/j.dadm.2016.12.011

14 Fernando, M. S. et al. Comparison of the pathology of cerebral white matter with post-mortem magnetic resonance imaging (MRI) in the elderly brain. Neuropathol. Appl. Neurobiol. 30, 385–395 (2004). 10.1111/j.1365-2990.2004.00550.x

15 Topakian, R., Barrick, T. R., Howe, F. A. & Markus, H. S. Blood-brain barrier permeability is increased in normal-appearing white matter in patients with lacunar stroke and leucoaraiosis. J. Neurol. Neurosurg. Psychiatry 81, 192–197 (2010). 10.1136/jnnp.2009.172072

16 Peters, A. The effects of normal aging on myelin and nerve fibers: a review. J. Neurocytol. 31, 581–593 (2002). 10.1023/a:1025731309829

17 Sargurupremraj, M. et al. Cerebral small vessel disease genomics and its implications across the lifespan. Nat. Commun. 11, 6285 (2020). 10.1038/s41467-020-19111-2

18 Moroni, F. et al. Cardiovascular disease and brain health: Focus on white matter hyperintensities. Int. J. Cardiol. Heart Vasc. 19, 63–69 (2018). 10.1016/j.ijcha.2018.04.006

19 Gorelick, P. B. et al. Vascular contributions to cognitive impairment and dementia: a statement for healthcare professionals from the american heart association/american stroke association. Stroke 42, 2672–2713 (2011). 10.1161/STR.0b013e3182299496

20 Jochems, A. C. C. et al. Longitudinal changes of white matter hyperintensities in sporadic small vessel disease. Neurology 99 (2022). 10.1212/wnl.0000000000201205

21 Duering, M. et al. Neuroimaging standards for research into small vessel disease-advances since 2013. Lancet Neurol. 22, 602–618 (2023). 10.1016/S1474-4422(23)00131-X

22 Scott, J. A. et al. Cerebral amyloid is associated with greater white-matter hyperintensity accrual in cognitively normal older adults. Neurobiol. Aging 48, 48–52 (2016). 10.1016/j.neurobiolaging.2016.08.014

23 Alosco, M. L. et al. A clinicopathological investigation of white matter hyperintensities and Alzheimer’s disease neuropathology. J. Alzheimers. Dis. 63, 1347–1360 (2018). 10.3233/JAD-180017

24 Kantarci, K. et al. White-matter integrity on DTI and the pathologic staging of Alzheimer’s disease. Neurobiol. Aging 56, 172–179 (2017). 10.1016/j.neurobiolaging.2017.04.024

25 McAleese, K. E. et al. Frontal white matter lesions in Alzheimer’s disease are associated with both small vessel disease and AD-associated cortical pathology. Acta Neuropathol. 142, 937–950 (2021). 10.1007/s00401-021-02376-2

26 McAleese, K. E. et al. Parietal white matter lesions in Alzheimer’s disease are associated with cortical neurodegenerative pathology, but not with small vessel disease. Acta Neuropathol. 134, 459–473 (2017). 10.1007/s00401-017-1738-2

27 Hyman, B. T. et al. National Institute on Aging–Alzheimer’s Association guidelines for the neuropathologic assessment of Alzheimer’s disease. Alzheimers. Dement. 8, 1–13 (2012). 10.1016/j.jalz.2011.10.007

28 Cerri, S. et al. A contrast-adaptive method for simultaneous whole-brain and lesion segmentation in multiple sclerosis. Neuroimage 225, 117471 (2021). 10.1016/j.neuroimage.2020.117471

29 Puonti, O., Iglesias, J. E. & Van Leemput, K. Fast and sequence-adaptive whole-brain segmentation using parametric Bayesian modeling. Neuroimage 143, 235–249 (2016). 10.1016/j.neuroimage.2016.09.011

30 Bryant, A. et al. Endothelial cells are heterogeneous in different brain regions and are dramatically altered in Alzheimer’s disease. J. Neurosci. 43, 4541–4557 (2023). 10.1523/JNEUROSCI.0237-23.2023

31 Holland, C. M. et al. Spatial distribution of white-matter hyperintensities in Alzheimer disease, cerebral amyloid angiopathy, and healthy aging. Stroke 39, 1127–1133 (2008). 10.1161/STROKEAHA.107.497438

32 Bryant, A. G. et al. Cerebrovascular senescence is associated with tau pathology in Alzheimer’s Disease. Front. Neurol. 11, 575953 (2020). 10.3389/fneur.2020.575953

33 Hoglund, Z. et al. Brain vasculature accumulates tau and is spatially related to tau tangle pathology in Alzheimer’s disease. bioRxiv (2024). 10.1101/2024.01.27.577088

34 Soneson, C., Love, M. I. & Robinson, M. D. Differential analyses for RNA-seq: transcript-level estimates improve gene-level inferences. F1000Res. 4, 1521 (2015). 10.12688/f1000research.7563.2

35 Soneson, C., Matthes, K. L., Nowicka, M., Law, C. W. & Robinson, M. D. Isoform prefiltering improves performance of count-based methods for analysis of differential transcript usage. Genome Biol. 17, 12 (2016). 10.1186/s13059-015-0862-3

36 Love, M. I., Huber, W. & Anders, S. Moderated estimation of fold change and dispersion for RNA-seq data with DESeq2. Genome Biol. 15, 550 (2014). 10.1186/s13059-014-0550-8

37 Zhou, Y. et al. Metascape provides a biologist-oriented resource for the analysis of systems-level datasets. Nat. Commun. 10, 1523 (2019). 10.1038/s41467-019-09234-6

38 Ge, S. X., Jung, D. & Yao, R. ShinyGO: a graphical gene-set enrichment tool for animals and plants. Bioinformatics 36, 2628–2629 (2020). 10.1093/bioinformatics/btz931

39 Koopmans, F. et al. SynGO: An evidence-based, expert-curated knowledge base for the synapse. Neuron 103, 217-234.e214 (2019). 10.1016/j.neuron.2019.05.002

40 Sakabe, M. et al. YAP/TAZ-CDC42 signaling regulates vascular tip cell migration. Proc. Natl. Acad. Sci. U. S. A. 114, 10918–10923 (2017). 10.1073/pnas.1704030114

41 Vabulas, R. M., Raychaudhuri, S., Hayer-Hartl, M. & Hartl, F. U. Protein folding in the cytoplasm and the heat shock response. Cold Spring Harb. Perspect. Biol. 2, a004390 (2010). 10.1101/cshperspect.a004390

42 Houry, W. A. Chaperone-assisted protein folding in the cell cytoplasm. Curr. Protein Pept. Sci. 2, 227–244 (2001). 10.2174/1389203013381134

43 Fazekas, F., Chawluk, J. B., Alavi, A., Hurtig, H. I. & Zimmerman, R. A. MR signal abnormalities at 1.5 T in Alzheimer’s dementia and normal aging. AJR Am. J. Roentgenol. 149, 351–356 (1987). 10.2214/ajr.149.2.351

44 Ylikoski, A. et al. White matter hyperintensities on MRI in the neurologically nondiseased elderly. Analysis of cohorts of consecutive subjects aged 55 to 85 years living at home. Stroke 26, 1171–1177 (1995). 10.1161/01.str.26.7.1171

45 Arana, M. E., Seki, M., Wood, R. D., Rogozin, I. B. & Kunkel, T. A. Low-fidelity DNA synthesis by human DNA polymerase theta. Nucleic Acids Res. 36, 3847–3856 (2008). 10.1093/nar/gkn310

46 Hull, J. et al. Distribution of the branched chain aminotransferase proteins in the human brain and their role in glutamate regulation. J. Neurochem. 123, 997–1009 (2012). 10.1111/jnc.12044

47 Ashby, E. L. et al. Altered expression of human mitochondrial branched chain aminotransferase in dementia with Lewy bodies and vascular dementia. Neurochem. Res. 42, 306–319 (2017). 10.1007/s11064-016-1855-7

48 Tubi, M. A. et al. White matter hyperintensities and their relationship to cognition: Effects of segmentation algorithm. Neuroimage 206, 116327 (2020). 10.1016/j.neuroimage.2019.116327

49 Luo, J. et al. Longitudinal relationships of white matter hyperintensities and Alzheimer disease biomarkers across the adult life span. Neurology 101, e164–e177 (2023). 10.1212/WNL.0000000000207378

50 Gordon, B. A. et al. The effects of white matter hyperintensities and amyloid deposition on Alzheimer dementia. NeuroImage Clin. 8, 246–252 (2015). 10.1016/j.nicl.2015.04.017

51 Brickman, A. M., Muraskin, J. & Zimmerman, M. E. Structural neuroimaging in Altheimer’s disease: do white matter hyperintensities matter? Dialogues Clin. Neurosci. 11, 181–190 (2009). 10.31887/dcns.2009.11.2/ambrickman

52 Debette, S. & Markus, H. S. The clinical importance of white matter hyperintensities on brain magnetic resonance imaging: systematic review and meta-analysis. BMJ 341, c3666 (2010). 10.1136/bmj.c3666

53 Gronewold, J. et al. Association of regional white matter hyperintensities with hypertension and cognition in the population-based 1000BRAINS study. Eur. J. Neurol. (2023). 10.1111/ene.15716

54 Marquine, M. J. et al. Differential patterns of cognitive decline in anterior and posterior white matter hyperintensity progression. Stroke 41, 1946–1950 (2010). 10.1161/STROKEAHA.110.587717

55 Salvadó, G. et al. Spatial patterns of white matter hyperintensities associated with Alzheimer’s disease risk factors in a cognitively healthy middle-aged cohort. Alzheimers. Res. Ther. 11, 12 (2019). 10.1186/s13195-018-0460-1

56 Alix, J. J. P. & de Jesus Domingues, A.M. White matter synapses. Neurology 76, 397–404 (2011). 10.1212/wnl.0b013e3182088273

57 Bakiri, Y. et al. Glutamatergic signaling in the brain’s white matter. Neuroscience 158, 266–274 (2009). 10.1016/j.neuroscience.2008.01.015

58 Sóki, N. et al. Investigation of synapses in the cortical white matter in human temporal lobe epilepsy. Brain Res. 1779, 147787 (2022). 10.1016/j.brainres.2022.147787

59 Patterson, E. K. et al. Proteinase 3 contributes to endothelial dysfunction in an experimental model of sepsis. Exp. Biol. Med. (Maywood) 246, 2338–2345 (2021). 10.1177/15353702211029284

60 Song, Y.-H., Yoon, J. & Lee, S.-H. The role of neuropeptide somatostatin in the brain and its application in treating neurological disorders. Exp. Mol. Med. 53, 328–338 (2021). 10.1038/s12276-021-00580-4

61 Chen, S., Chen, H., Du, Q. & Shen, J. Targeting myeloperoxidase (MPO) mediated oxidative stress and inflammation for reducing brain ischemia injury: Potential application of natural compounds. Front. Physiol. 11, 433 (2020). 10.3389/fphys.2020.00433

62 Zhao, H. et al. Lipocalin 10 is essential for protection against inflammation-triggered vascular leakage by activating LRP2-Ssh1 signaling pathway. Cardiovasc. Res. (2023). 10.1093/cvr/cvad105

63 Liu, B. et al. Protective effect of KLF15 on vascular endothelial dysfunction induced by TNF-α. Mol. Med. Rep. 18, 1987–1994 (2018). 10.3892/mmr.2018.9195

64 Jickling, G. C. et al. Progression of cerebral white matter hyperintensities is related to leucocyte gene expression. Brain 145, 3179–3186 (2022). 10.1093/brain/awac107

65 Sole-Guardia, G. et al. Three-dimensional identification of microvascular pathology and neurovascular inflammation in severe white matter hyperintensity: a case report. Sci Rep 14, 5004 (2024). 10.1038/s41598-024-55733-y

66 Yang, Y. et al. Epigenetic and integrative cross-omics analyses of cerebral white matter hyperintensities on MRI. Brain 146, 492–506 (2023). 10.1093/brain/awac290

67 Noz, M. P. et al. Pro-inflammatory monocyte phenotype during acute progression of cerebral small vessel disease. Front. Cardiovasc. Med. 8, 639361 (2021). 10.3389/fcvm.2021.639361

68 Lin, H. et al. Whole blood gene expression and white matter Hyperintensities. Mol. Neurodegener. 12 (2017). 10.1186/s13024-017-0209-5

69 Raina, A. et al. Cerebral white matter hyperintensities on MRI and acceleration of epigenetic aging: the atherosclerosis risk in communities study. Clin. Epigenetics 9, 21 (2017). 10.1186/s13148-016-0302-6

70 Farkhondeh, V. & DeCarli, C. White matter hyperintensities in diverse populations: A systematic review of literature in the United States. Cereb Circ Cogn Behav 6, 100204 (2024). 10.1016/j.cccb.2024.100204

71 Sachdev, P., Wen, W., Chen, X. & Brodaty, H. Progression of white matter hyperintensities in elderly individuals over 3 years. Neurology 68, 214–222 (2007). 10.1212/01.wnl.0000251302.55202.73

72 Maillard, P. et al. White matter hyperintensities and their penumbra lie along a continuum of injury in the aging brain. Stroke 45, 1721–1726 (2014). 10.1161/STROKEAHA.113.004084

