## Supplementary Information and Figures for "Molecular profiling of frontal and occipital subcortical white matter hyperintensities in Alzheimer’s disease"

**Supplementary Tables**

Sup. Table 1. Differentially Expressed Genes (DEGs) in Frontal-WM bulk tissue (2198 DEGs; 1033 up; 1165 down)

Sup. Table 2. Biological processes enriched in significantly up- or downregulated DEGs in Frontal-WM bulk tissues

Sup. Table 3. Differential Expressed Genes (DEGs) in Occipital-WM-bulk tissue (208 DEGs; 155 up; 53 down)

Sup. Table 4. Biological processes enriched in significantly up- or downregulated DEGs in Occipital-WM bulk tissues

Sup. Table 5. Differentially Expressed Genes (DEGs) in Frontal-WM blood vessels (690 DEGs; 568 up; 122 down)

Sup. Table 6. Biological processes enriched in significantly up- or downregulated DEGs in Frontal-WM blood vessels

Sup. Table 7. Differentially Expressed Genes (DEGs) in Occipital-WM blood vessels (133 DEGs; 70 up; 63 down)

Sup. Table 8. Biological processes enriched in significantly up- or downregulated DEGs in Occipital-WM blood vessels

Sup. Table 9. High WMH vs. Low WMH Differentially Expressed Genes (DEGs) in Frontal-WM Bulk Tissue

Sup. Table 10. High WMH vs. Low WMH Differentially Expressed Genes (DEGs) in Frontal-WM Blood Vessels

Sup. Table 11. High WMH vs. Low WMH Differentially Expressed Genes (DEGs) in Occipital-WM Bulk Tissue

Sup. Table 12. High WMH vs. Low WMH Differentially Expressed Genes (DEGs) in Occipital-WM Blood Vessels

Sup. Table 13. Biological Processes enriched in significantly upregulated genes in high vs. low WMH occipital-WM bulk tissues

Sup. Table. 14. Biological processes enriched in significantly upregulated genes in high- vs. low. WMH occipital-WM blood vessels

### Supplementary Figures

A

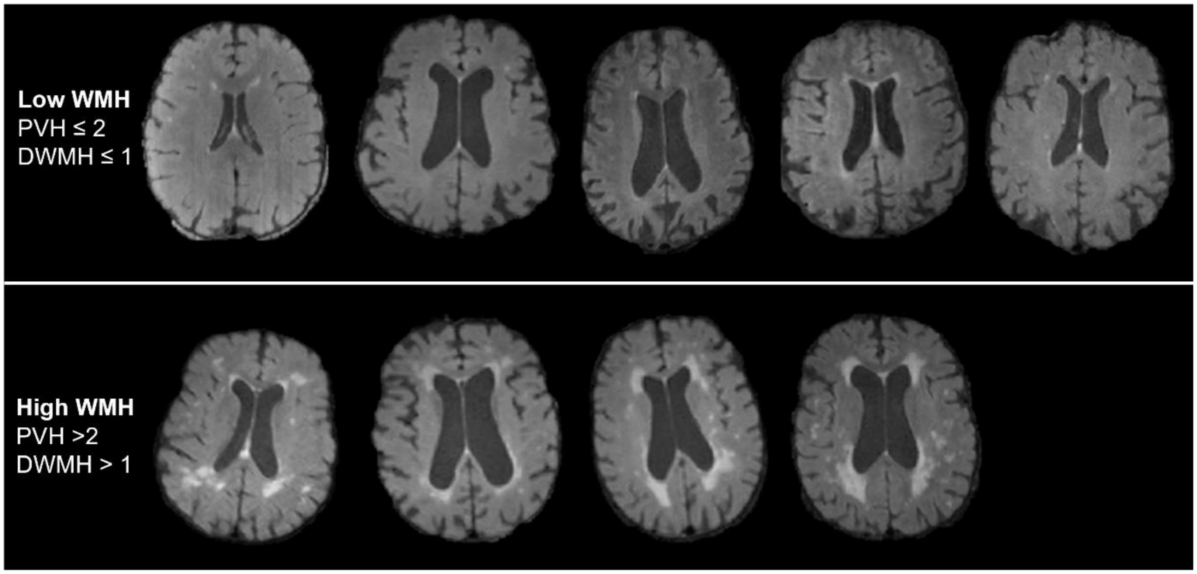


B

| Group | Age (Mean ± STD) | Braak stage | Thal stage | Avg. pre-mortem scan int. |
|---|---|---|---|---|
| **Control (n=5)** | 85 ± 5.1 | I-II | 0-3 | NA |
| **AD, Low WMH (n=5)** | 77 ± 8.8 | V-VI | 4-5 | 2.95 years |
| **AD, High WMH (n=4)** | 85 ± 5.4 | V-VI | 4-5 | 2.81 years |

**Supplementary Figure 1. A)** *In-vivo* T2-weighted FLAIR MRI showing the low (upper panel) and high (lower panel) WMH burden AD in donors (PVH- periventricular hyperintensities; DWMH = deep white matter hyperintensities). **B**) Information showing group averages for age, AD pathology (Braak and Thal staging), and pre-mortem scan interval.

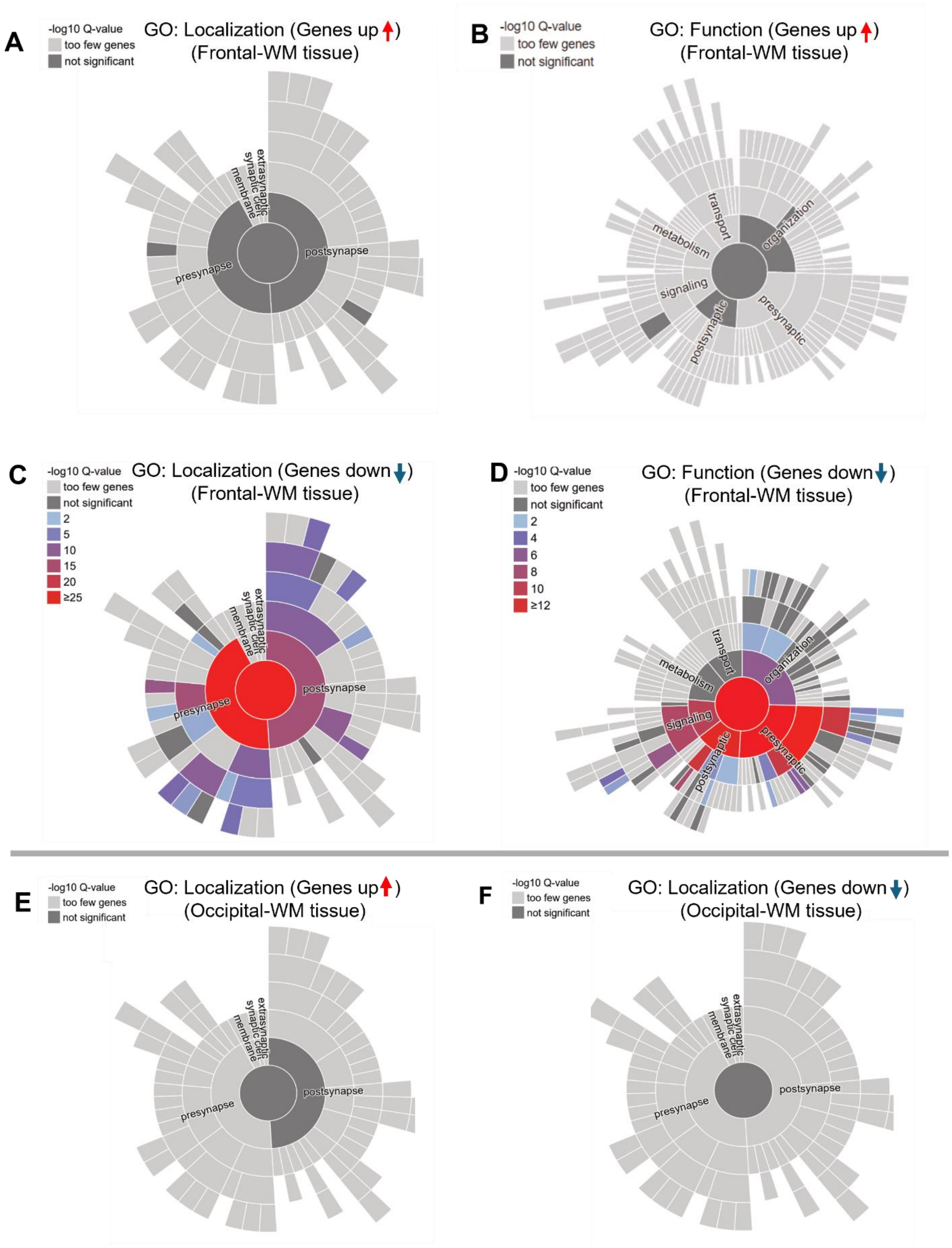


**Supplementary Figure 2. A-B)** Sunburst image of upregulated genes in frontal-WM bulk tissues in control versus AD with WMH, generated by SynGO showing **A)** cellular component enrichment, and (**B)** functional enrichment. **C-D**) Same as A & B but for downregulated genes in frontal-WM bulk tissues. **E-F**) Sunburst image showing the cellular component enrichment of (**E**) upregulated and (**F**) downregulated genes in occipital-WM bulk tissues. The red color in the center of the sunburst plot indicates the higher enrichment of significant synapse related genes while the light gray color suggests only a few genes are enriched in synaptic cellular component and dark gray color in the center of the sunburst plot indicates no significant enrichment of synapse related genes. The upward and downward pointed arrows refer to upregulated and downregulated genes, respectively.
